# Multiple origins, one function: evolutionary pathways of HSP70 proteins in viruses

**DOI:** 10.1101/2025.07.06.663342

**Authors:** Ayoub Maachi, Santiago F. Elena

## Abstract

Heat shock proteins 70 (HSP70) are highly conserved molecular chaperones found across all domains of life, where they play essential roles in cellular stress responses. While HSP70 homologs have been previously identified in closteroviruses (ssRNA viruses), their broader presence and evolutionary history in viruses remain poorly understood. In this study, we conducted a comprehensive search of viral protein databases and identified HSP70 homologs in viruses beyond ssRNA, including dsDNA viruses from the classes *Megaviricetes* and *Caudoviricetes*. These viral HSP70s exhibit diverse gene organizations, copy numbers, and structural features. Notably, HSP70s from *Megaviricetes* showed up to three gene copies per genome and distinct structural motifs, while those from closteroviruses displayed higher sequence and structural diversity, suggesting faster evolutionary rates. Structural and phylogenetic analyses revealed two major clusters of viral HSP70s, with dsDNA virus HSP70s closely resembling those of their protist hosts, supporting the hypothesis of horizontal gene transfer. In contrast, ssRNA virus HSP70s formed a distinct, highly divergent group. Our findings suggest multiple independent acquisitions of HSP70 genes by viruses and provide new insights into their evolutionary trajectories and potential functional adaptations.

## Introduction

Heat shock proteins (HSPs) are a highly conserved family of molecular chaperones found across all domains of life, including bacteria, archaea and eukaryotes (Hu et al., 2022). These proteins are essential for maintaining cellular proteostasis, particularly under conditions of physiological stress such as heat, oxidative damage, or infection. Among the various HSP families, the 70-kDa heat shock proteins (HSP70s) are among the most studied due to their central role in protein folding, assembly, translocation, and degradation (Rosenzweig et al., 2019). Structurally, HSP70s are composed of a nucleotide-binding domain (NBD) and a substrate-binding domain (SBD), connected by a flexible linker. The SBD itself includes a substrate-binding pocket and a lid domain, often ending with a con served EEVD motif that mediates interactions with co-chaperones (Fernández-Fernández et al., 2017; Zhang & Zuiderweg, 2004). Large HSPs such as HSP110 and Grp170, which are homologous to HSP70s, also contribute to protein quality control and are considered part of the HSP70 superfamily (Easton et al., 2000; Raviol et al., 2006; Tittelmeier et al., 2020).

In the context of viral infections, HSP70s play dual roles. On the one hand, they can inhibit viral replication by interfering with viral protein function or stability (Wang et al., 2020). On the other hand, viruses can exploit host HSP70s to facilitate their own entry, replication, and assembly. For example, HSP70s have been shown to act as viral receptors (Chuang et al., 2015), assist in membrane translocation (Ravindran et al., 2015), stabilize viral ribonucleoproteins (Naito et al., 2007), and sup-port virion assembly (Gurer et al., 2005). Despite their functional importance during infection, relatively little is known about whether viruses themselves encode HSP70 homologs. To date, the best-known examples are found in the plant-infecting ssRNA closteroviruses, which encode HSP70-like proteins believed to have been acquired from host mRNAs via recombination (Agranovsky et al., 1991; Dolja et al., 1994). These viral HSP70s retain conserved ATPase domains but show divergence in their C-terminal regions, and they function in viral movement between plant cells (Peremyslov et al., 1999).

In this study, we aimed to systematically investigate the presence, diversity, and evolutionary ori gins of HSP70 proteins encoded by viruses. We conducted a comprehensive search of viral protein sequences in the NCBI database, identifying HSP70 homologs in both ssRNA and dsDNA viruses, including members of the *Alsuviricetes*, *Caudoviricetes* and *Megaviricetes* classes. We analyzed their gene organization, structural features, sequence diversity, phylogenetic relationships, and possible functional diversification. Our findings reveal multiple independent acquisitions of HSP70 genes by viruses, distinct evolutionary trajectories between RNA and DNA viruses, and potential horizontal gene transfer (HGT) events from host organisms, particularly protists. This work provides novel insights into the evolutionary plasticity and functional adaptation of viral genomes.

## Results

### Distribution and abundance of viral HSP70s

We retrieved 63 viral HSP70 sequences from the NCBI protein database, with lengths ranging from 533 to 1147 amino acids (Figure 1, Table S1). These sequences were primarily from two viral classes: *Alsuviricetes* (ssRNA viruses infecting plants) and *Megaviricetes* (giant dsDNA viruses infecting protists). *Alsuviricetes*, specifically members of the *Closteroviridae* family, accounted for 68% of the sequences, with HSP70 lengths between 533 and 606 amino acids. *Megaviricetes* included viruses from the orders *Algavirales* and *Imitervirales*, and families *Phycodnaviridae* and *Mimiviridae*, respectively. *Phycodnaviruses* (8%) infect algae, while mimiviruses (20%) infect amoebae.

**Figure 1.**
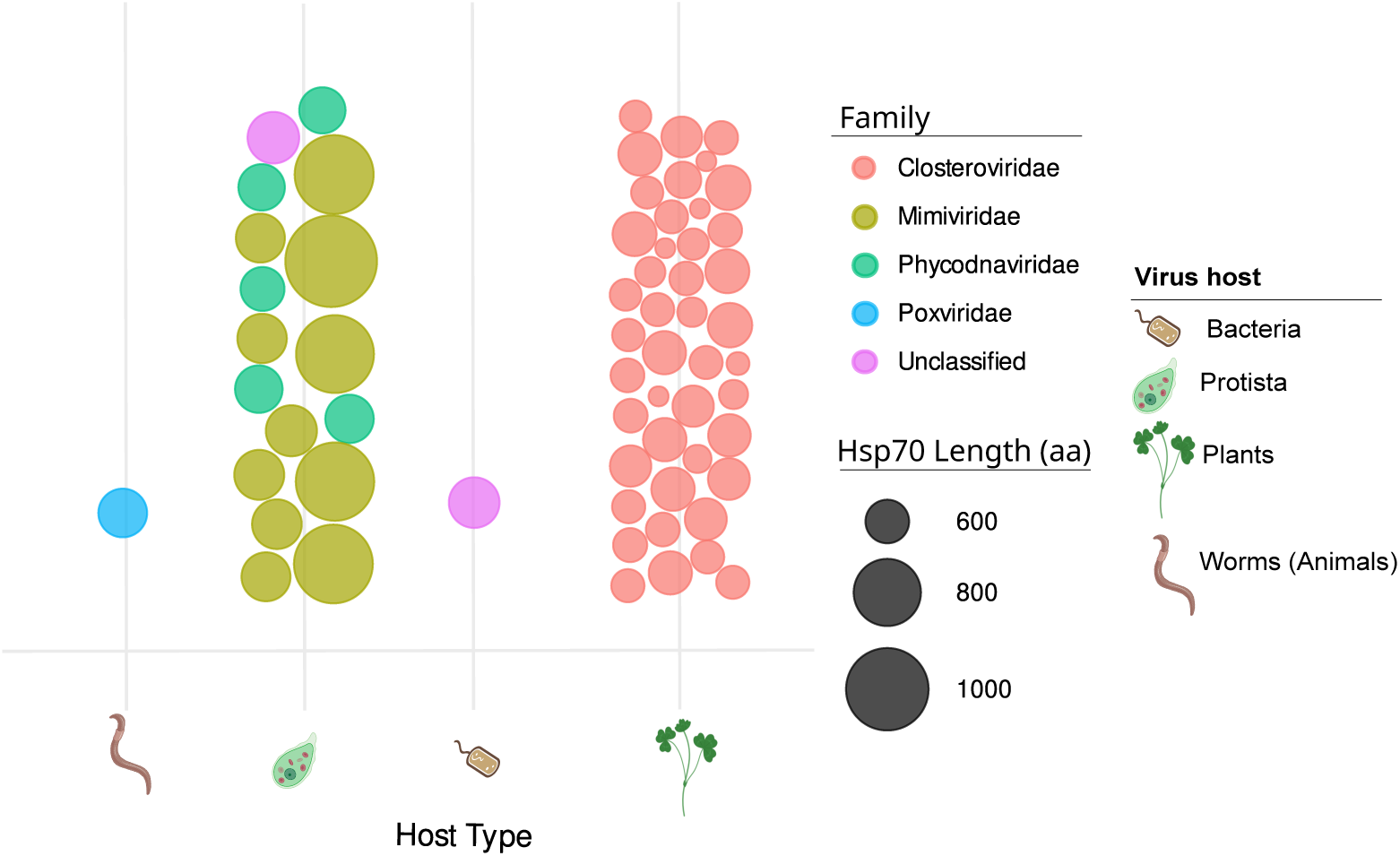
Abundance of heat shock protein 70 (HSP70) homologs in viruses, categorized by protein size, host organism, and viral family.

Two additional sequences were identified from less-represented classes: an *Acinetobacter phage* (CAH91635, *Caudoviricetes*) and *Poxvirus euperiparoides* (DBA47115, *Pokkesviricetes*), infecting bacteria and velvet worms, respectively. BLASTp analysis for the *Acinetobacter* phage HSP70 yielded homologs to the DnaK chaperone from 25 related bacteria (> 96% identity), while no additional homologs were found for the poxvirus sequence, suggesting a possible misannotation.

### Copy number variation in *Megaviricetes*

Among *Imitervirales* members, several viruses encoded multiple HSP70 copies. *Acanthamoeba castellanii mimivirus*, *Catovirus naegleriensis*, and *Cotonvirus japonicus* each encoded two HSP70s (∼600 and ∼900 amino acids). Other viruses, such as *Bandra megavirus*, *Hyperinonvirus*, and *Yas-minevirus*, encoded single large HSP70s (> 900 amino acids) (Table S1).

To investigate this further, we analyzed complete genomes of *Imitervirales*. Most encoded two HSP70s, except *C. naegleriensis* and *Yasminevirus*, which had three (Figure 2A, Table S2). These genes were randomly distributed across both DNA strands and showed no correlation with protein length. Pairwise amino acid identity ranged from 39% to 88%, with small HSP70s (< 655 amino acids) showing higher similarity (> 80%) among *Moumouvirus australiensis*, *Megavirus courdo 11* and *A. castellanii mimivirus* (Figure S1).

**Figure 2.**
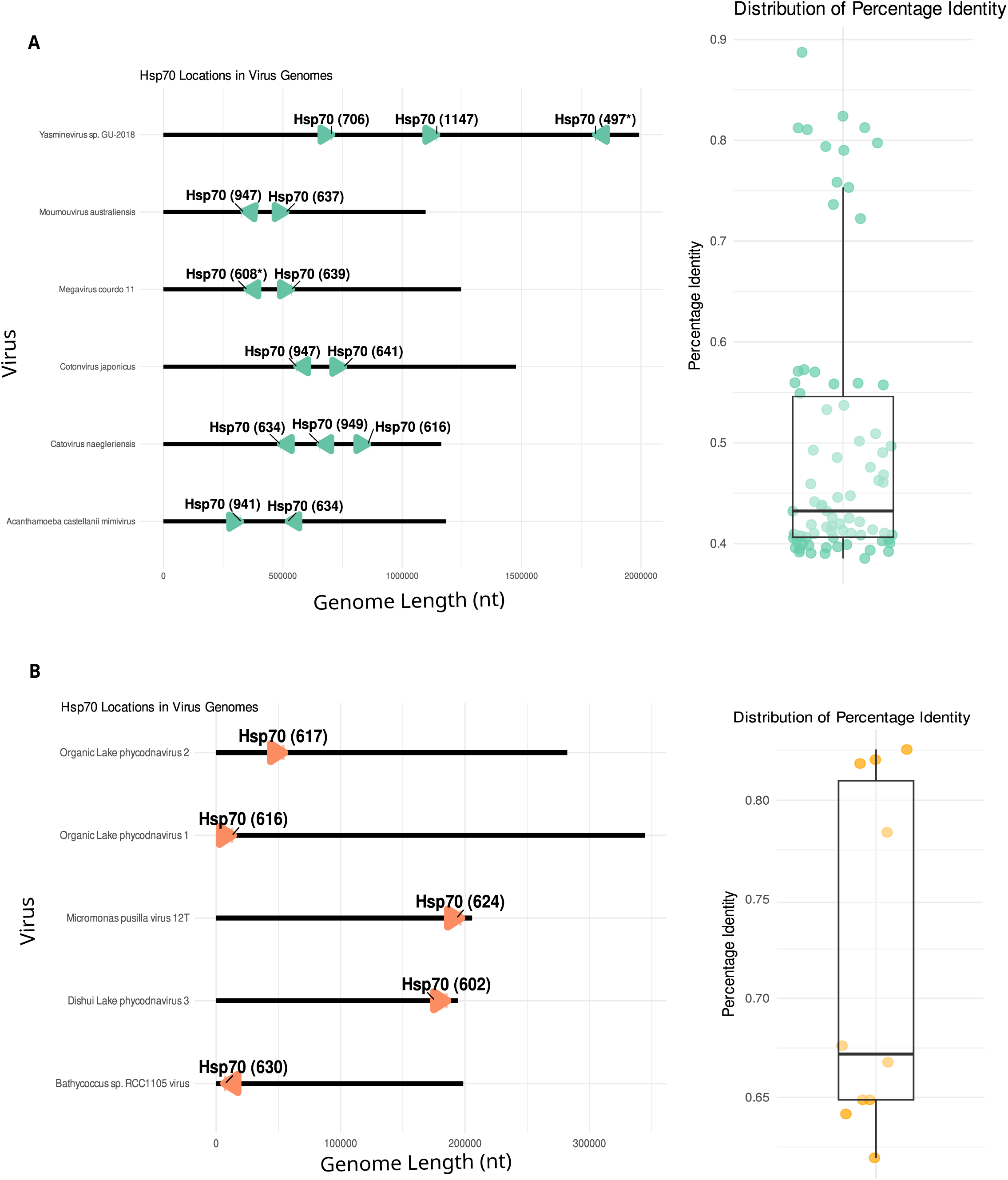
Genomic organization of HSP70 genes in viruses. (A) *Imitervirales* members: gene location, length, strand orientation, and pairwise identity. (B) *Algavirales* members: similar features as in (A). Fragmented HSP70s are marked with an asterisk (*).

In contrast, *Algavirales* members encoded only one HSP70 per genome, all small (602 - 624 amino acids), located primarily on the sense strand (Figure 2B, Table S3). Pairwise identity among these proteins ranged from 61% to 81% (Figure 2B).

### Gene structure and expression patterns

Most viral HSP70s were encoded by single-exon genes. However, exceptions were observed in *M. courdo 11* and *Yasminevirus*. In *M. courdo 11*, one HSP70 gene (639 amino acids) contained an intron, while another (609 amino acids) appeared split across two intergenic regions, producing two immature proteins that aligned structurally to form a complete HSP70 (Figure S2A - B). Similarly, *Yas-minevirus* encoded two gene fragments separated by an unrelated ORF, which together formed a full-length HSP70 (Figure S2C). These findings suggest alternative splicing or gene fragmentation mechanisms in some *Imitervirales*.

### Structural and motif diversity

To assess structural diversity, we predicted 3D models of all viral HSP70s and performed correspondence analysis (CA) based on structural similarity. The CA revealed two major clusters (Figure 3A): one comprising *Closteroviridae* (ssRNA viruses) with high structural dispersion, and another comprising dsDNA viruses with more compact structures. The main structural differences were observed in the substrate-binding domain (SBD), which was shorter and more variable in ssRNA viruses (Figure 3B).

**Figure 3.**
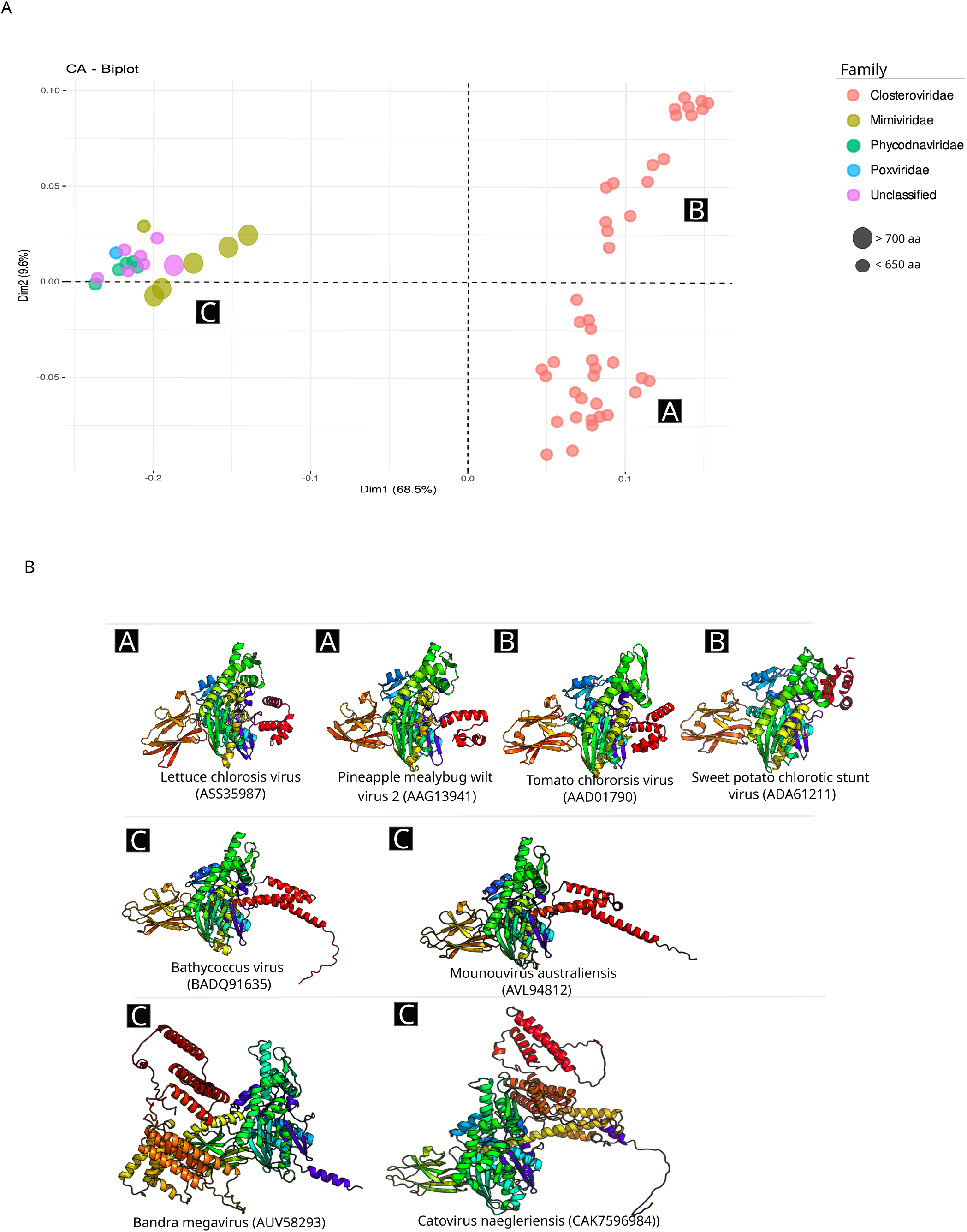
Structural diversity of viral HSP70 proteins. (A) Correspondence analysis (CA) based on structural similarity, with proteins labeled by virus family and size. (B) Representative 3D structures from each CA cluster, colored from N-terminus (blue) to C-terminus (red).

Motif analysis identified eight conserved motifs (1 - 8) present in all viral HSP70s, primarily located in the NBD (Figure 4B, Table S4). Four additional motifs (9, 10, 13, and 17) were specific to dsDNA viruses, located in both the SBD and NBD (Figure 5A). Conversely, four motifs unique to *Closteroviridae* were longer (26 - 50 amino acids) and variably distributed (Figure 5B). Notably, all *Closteroviridae* HSP70s lacked the canonical EEVD motif at the C-terminus.

**Figure 4.**
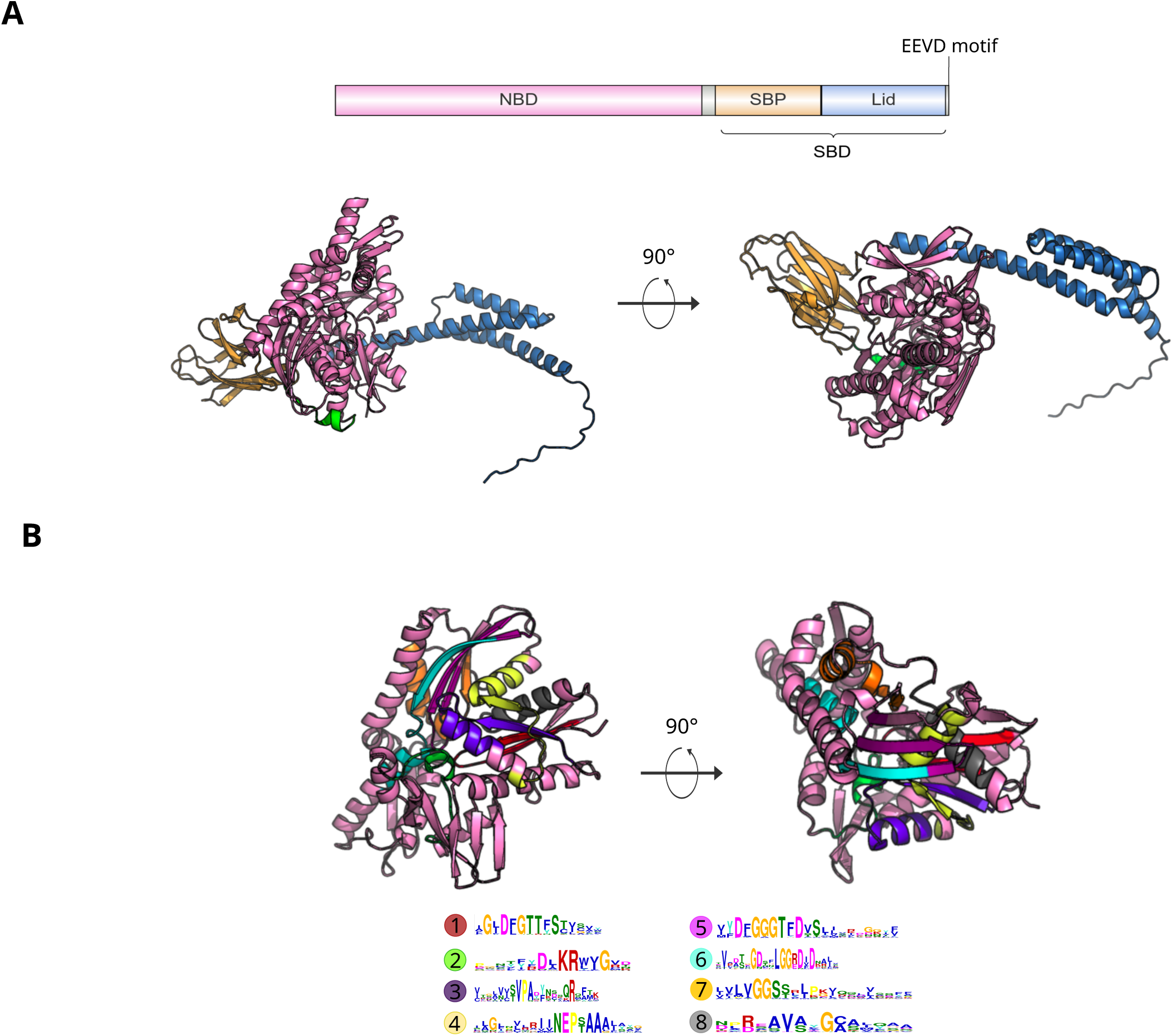
Domain architecture and conserved motifs in viral HSP70s. (A) Schematic of HSP70 domains from *Bathycoccus virus* (accession: BADQ91635), showing the nucleotide-binding domain (NBD) and substrate-binding domain (SBD), including the lid and substrate-binding pocket (SBP). (B) Positions of conserved motifs (1 - 8) within the NBD across all viral HSP70s. Full motif sequences are listed in Table S4.

**Figure 5.**
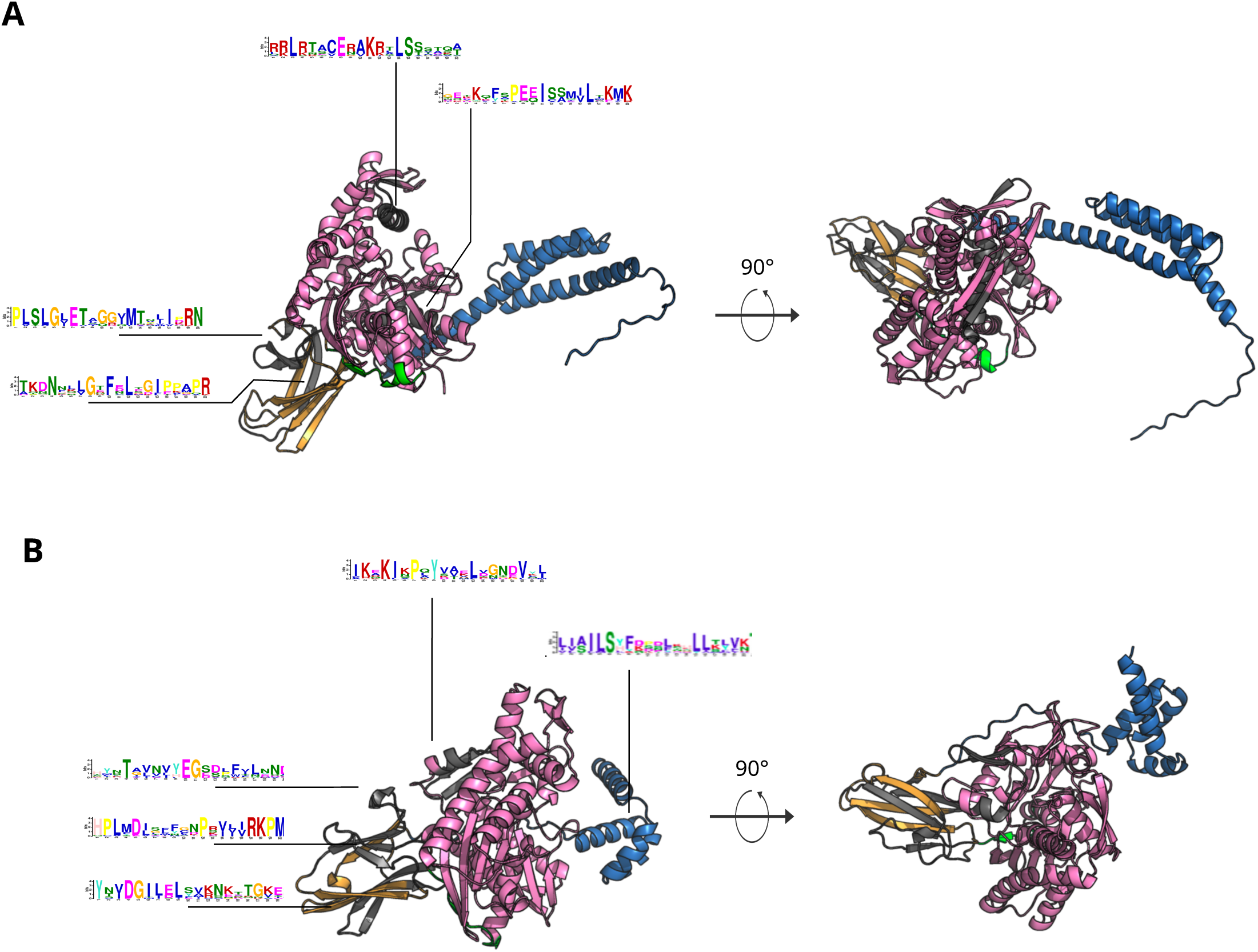
Lineage-specific motifs in viral HSP70s. (A) Motifs unique to dsDNA viruses (giant viruses), mapped onto the HSP70 from Bathycoccus virus (BADQ91635). (B) Motifs specific to ss-RNA viruses (closteroviruses), illustrated using the HSP70 from *Areca palm velarivirus 1* (YP_009140434).

### Phylogenetic relationships and host associations

Phylogenetic analysis of viral HSP70s revealed distinct clades (Figure 6). *Closteroviridae* formed a well-supported, diverse clade with long branch lengths, indicating high evolutionary rates. *Megaviricetes* sequences formed three groups: one of large mimiviral HSP70s (900 - 1100 amino acids), a second of smaller mimiviral HSP70s (∼600 amino acids), and a third combining phycodnaviruses and the poxvirus sequence.

**Figure 6.**
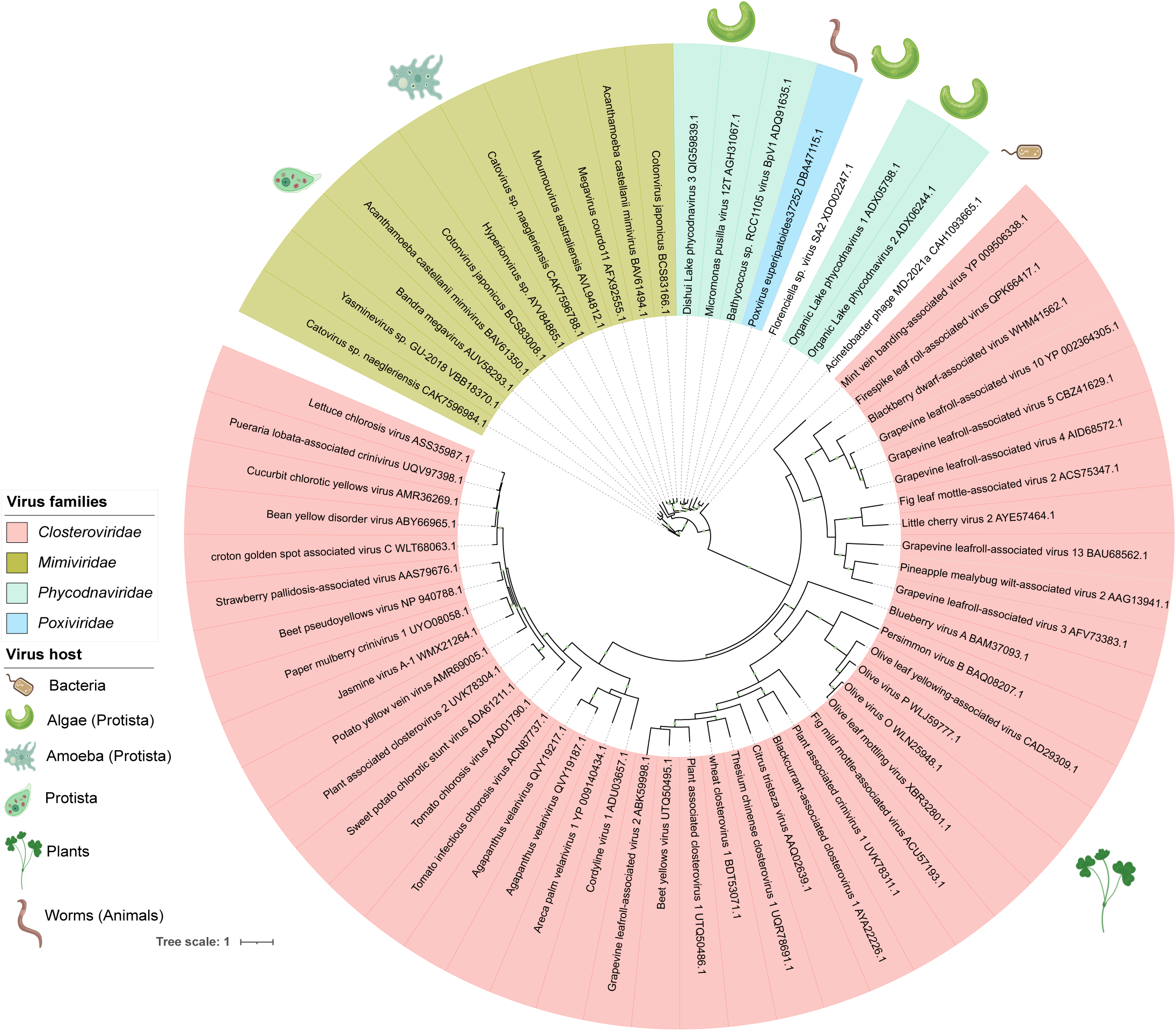
Phylogenetic tree of viral HSP70 proteins. Constructed using IQ-TREE with the LG+F+R5 substitution model. Bootstrap values > 70% (1,000 replicates) are shown.

To explore host-virus relationships, we constructed a broader phylogeny including HSP70s from bacteria, archaea, fungi, protists, plants, and animals (Figure 7). Viral sequences did not form a mono phyletic group. *Closteroviridae* clustered separately from all cellular taxa, while *Megaviricetes* HSP70s grouped near protist sequences, suggesting HGT from host to virus.

**Figure 7.**
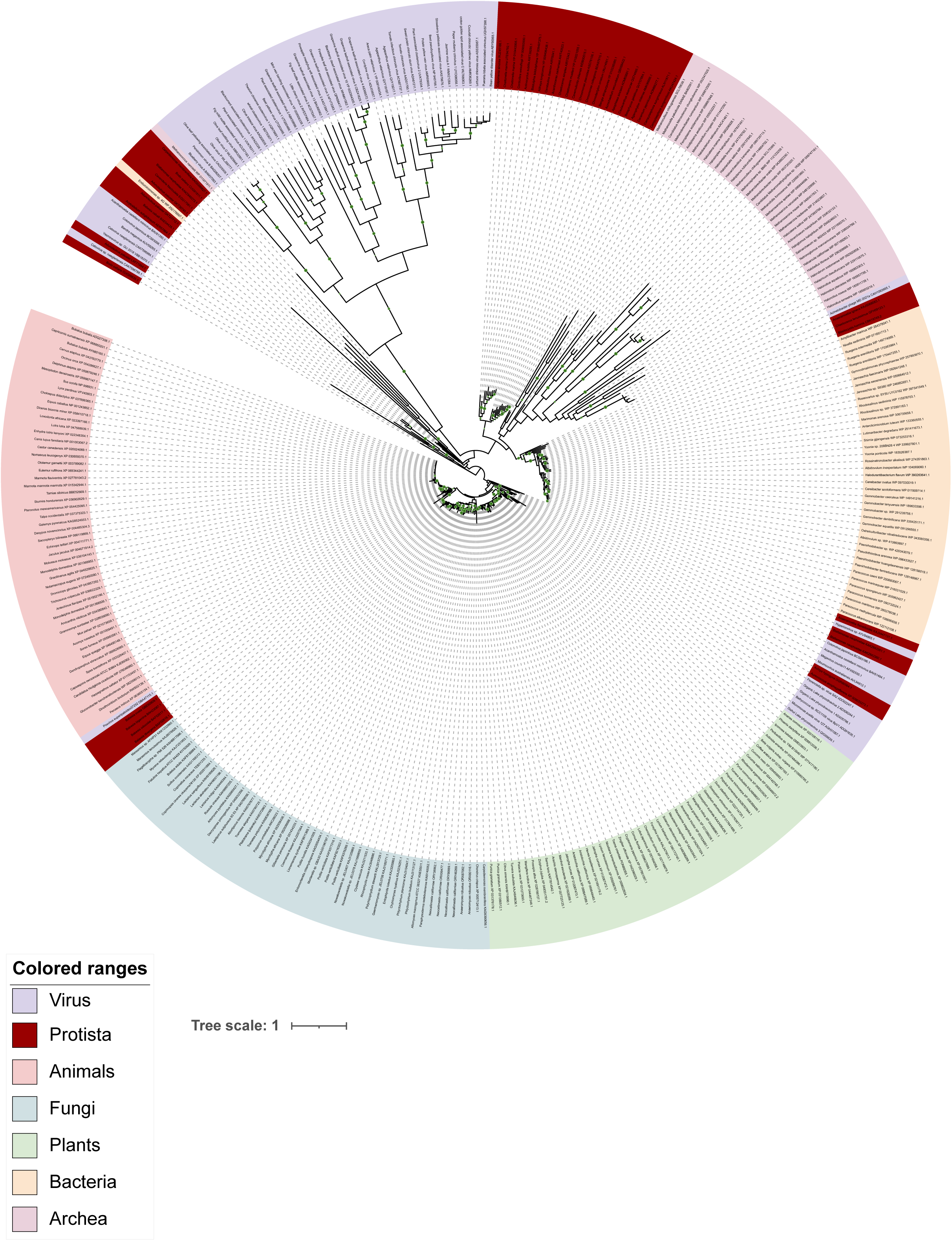
Phylogenetic relationships between viral and cellular HSP70s. Tree includes sequences from viruses, plants, fungi, protists, animals, and archaea. Constructed using RAxML with the PROTGAMMALG model. Bootstrap values > 70% (1,000 replicates) are shown.

### Functional network analysis reveals divergent evolutionary paths of viral HSP70s

To further explore the functional relationships among viral HSP70 proteins, we constructed sequence similarity networks using the EFI-Enzyme Similarity Tool (EFI-EST) (Oberg et al., 2023). At a strict alignment score threshold of 35, the network of viral HSP70s (Figure 8A) resolved into two completely disconnected clusters: one composed exclusively of HSP70s from ssRNA viruses (*Closteroviridae*), and the other comprising HSP70s from dsDNA viruses. This separation supports the hypothesis of distinct evolutionary origins for these groups. When the threshold was relaxed to 10 (Figure S3A), all viral HSP70s formed a single connected network; however, the spatial separation of nodes still indicated two major subgroups, consistent with functional divergence.

**Figure 8.**
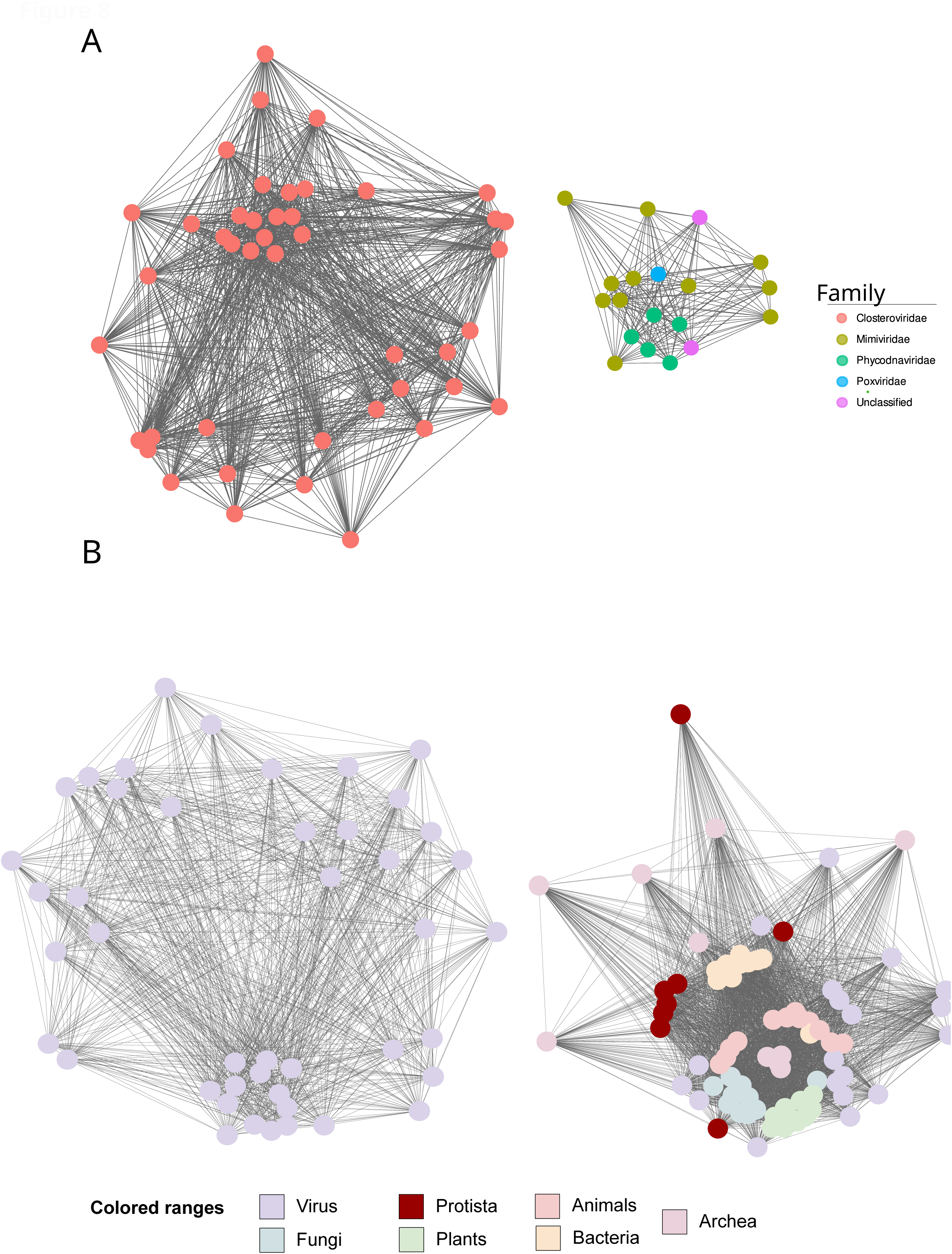
Sequence similarity network of viral and cellular HSP70 proteins generated using the EFI-EST tool. (A) Network of viral HSP70s constructed using a strict alignment score threshold of 35. Two completely disconnected clusters are observed: one composed of HSP70s from ssRNA viruses (*Closteroviridae*) and the other from dsRNA viruses, indicating strong sequence divergence. (B) Network including HSP70s from viruses and representative cellular organisms (10 species each from plants, animals, fungi, protists, bacteria, and archaea), constructed using the strict threshold (35). HSP70s from ssRNA viruses form a distinct, isolated cluster, while those from dsDNA viruses integrate with cellular HSP70s, mainly from protists.

To assess the relationship between viral and cellular HSP70s, we generated a network including representative sequences from major taxonomic groups (plants, animals, fungi, protists, bacteria, and archaea) along with all viral HSP70s, using the strict threshold (Figure 8B). In this network, HSP70s from ssRNA viruses again formed a separate, isolated cluster, while those from dsDNA viruses integrated with cellular sequences, particularly those from protists, further supporting HGT from host to virus. When the threshold was relaxed to 10 (Figure S3B), again all viral HSP70s formed a single con nected network; however, the spatial separation of nodes still indicated two major subgroups, consistent with functional divergence. These network-based findings reinforce our structural and phylogenetic analyses, highlighting both the evolutionary independence of ssRNA viral HSP70s and the host-derived origins of their dsDNA counterparts.

## Discussion

This study provides the first comprehensive analysis of heat shock protein 70 (HSP70) homologs encoded by viruses, revealing their limited but diverse presence across viral taxa and offering new insights into their structural features, gene organization, and evolutionary origins. Our results confirm that viral HSP70s are not widespread but are instead restricted to specific lineages, primarily ssRNA viruses from the *Closteroviridae* family and dsDNA viruses from *Mimiviridae* and *Phycodnaviridae*. This restricted distribution suggests that HSP70 acquisition is not a universal viral trait but rather a lineage-specific adaptation, likely driven by ecological or functional pressures. The presence of HSP70s in *Closteroviridae* has been previously reported (Agranovsky et al., 1991; Dolja et al., 1994), but our study expands this to include dsDNA viruses, particularly those infecting protists.

We observed that whilst most viruses encode a single HSP70 gene, members of *Imitervirales* often harbor two or three copies. These genes are randomly distributed across the genome and located on both strands, suggesting independent insertion events. In some cases, such as *M. courdo 11* and *Yasminevirus*, we identified atypical gene architectures, including intron-containing genes and fragmented open reading frames that still produce structurally coherent proteins, however, it is not clear whether these proteins retain their function, or if the loss of function has driven the virus to hijack other functional HSP70 from their hosts to fulfill that role. These findings suggest the presence of virus-specific mechanisms such as alternative splicing or post-translational assembly (Kovacs et al., 1991; Bouton et al., 2015; Dubois et al., 2014; Price et al., 2022), highlighting the remarkable genomic plasticity of giant viruses.

Structural modeling and correspondence analysis revealed two major HSP70 clusters: one comprising ssRNA viruses and another comprising dsDNA viruses. The ssRNA virus HSP70s exhibited greater structural variability, particularly in the substrate-binding domain (SBD), consistent with their higher evolutionary rates (Duffy et al., 2008). In contrast, dsDNA virus HSP70s were more structurally conserved. Motif analysis identified both universally conserved motifs and lineage-specific ones. Notably, all *Closteroviridae* HSP70s lacked the canonical EEVD motif, which is typically involved in co-chaperone interactions (Fernández-Fernández et al., 2017; Rosenzweig et al., 2019), suggesting functional divergence from their cellular counterparts.

Phylogenetic analyses revealed that viral HSP70s do not form a monophyletic group. Instead, they are scattered across the tree, with dsDNA virus HSP70s clustering near their protist hosts and ss RNA virus HSP70s forming a distinct, highly divergent clade. These patterns strongly suggest multiple independent HGT events from host organisms to viruses. The close relationship between *Megaviricetes* HSP70s and protist sequences supports the hypothesis of host-to-virus HGT (Zhang & Yu, 2022), while the long branch lengths of *Closteroviridae* HSP70s may reflect an ancient acquisition followed by rapid evolution (Dolja et al., 1994).

To further explore functional relationships among viral HSP70s, we constructed sequence similarity networks using the EFI-EST tool. At a strict alignment score threshold, viral HSP70s formed two completely disconnected clusters: one composed of plant ssRNA viruses and the other of dsDNA viruses. This separation reinforces the deep sequence divergence between these groups. When the threshold was relaxed, all viral HSP70s formed a single connected network, but the spatial separation of nodes still reflected two major subgroups, consistent with functional divergence. When cellular HSP70s were included in the network (using the strict threshold), ssRNA viral HSP70s again formed an isolated cluster, while dsDNA viral HSP70s integrated with cellular sequences, particularly from protists, further supporting the hypothesis of horizontal gene transfer from host to virus. These network-based findings align with our structural and phylogenetic results, providing additional evidence for the distinct evolutionary trajectories of viral HSP70s.

Based on our findings, we propose two evolutionary scenarios for dsDNA viruses: (*i*) a single ancestral acquisition of HSP70 in *Megaviricetes*, followed by divergence into *Algavirales* and *Imitervirales*, with the latter acquiring additional HSP70 copies (Figure 9A); or (*ii*) independent acquisitions in each lineage (Figure 9B). For ssRNA viruses, the data support a single, ancient acquisition from an unknown source, followed by extensive divergence (Figure 9C).

**Figure 9.**
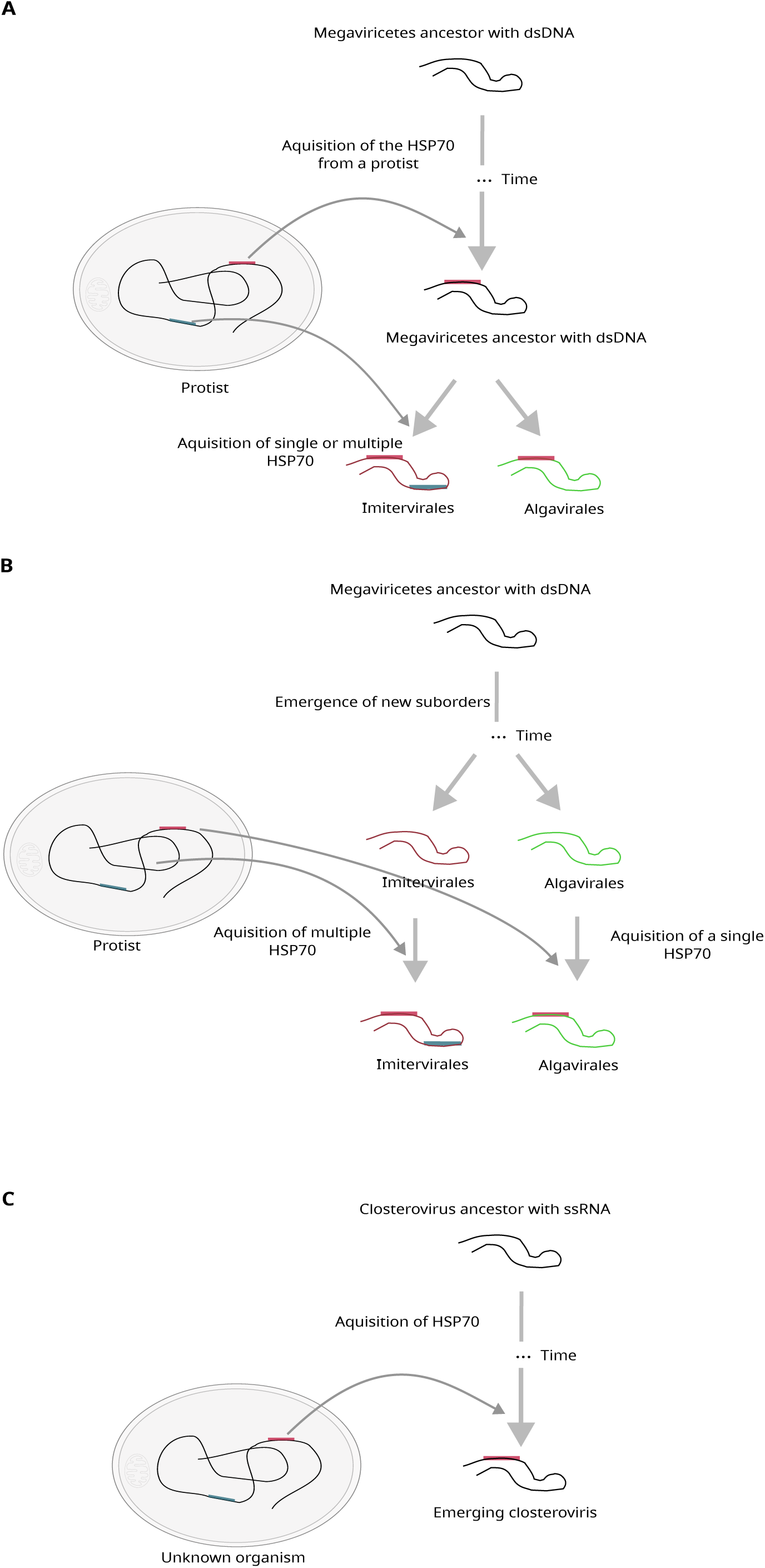
Proposed evolutionary scenarios for the acquisition of HSP70 genes in viruses. (A) Single acquisition in a *Megaviricetes* ancestor, followed by divergence into *Algavirales* and *Imitervirales*. (B) Independent acquisitions in each lineage. (C) Separate acquisition in closteroviruses from an unknown source.

This study provides foundational insights into the evolution and diversity of viral HSP70s. Future experimental work is needed to determine the functional roles of these proteins in viral replication, host interaction, and stress response, particularly in dsDNA viruses where their biological significance remains largely unexplored.

## Materials and Methods

### Retrieval of viral HSP70 sequences

A total of 74 viral HSP70 amino acid sequences were retrieved from the NCBI protein database using the search term “viruses” and excluding partial sequences. To avoid redundancy, only one sequence per species was retained, except for *C. japonicus*, *C. naegleriensis* and *A. castellanii mimivirus*, which each encode two distinct HSP70 proteins of different lengths. Sequences were clustered using CD-HIT with a 95% identity threshold (Li & Godzik, 2006), resulting in 63 representative sequences for downstream analyses (Table S1). Host information was obtained from the Virus-Host DB (https://www.genome.jp/virushostdb/view/) or the literature. Additionally, 50 HSP70 sequences from plants, animals, protists, fungi, archaea, and bacteria were randomly selected from NCBI for phylogenetic comparison.

### Identification of HSP70 homologs in giant viruses

To identify HSP70 homologs in giant viruses, we created a custom HSP70 database from the retrieved sequences and formatted it using the makeblastdb tool. All protein-coding sequences from each giant virus genome were downloaded and queried against the HSP70 database using BLASTp (Camacho et al., 2008). Hits with *E*-values ≤ 10⁻³ were retained and manually verified. When full or near-complete genomes were available, gene positions and orientations were annotated. Pairwise amino acid identities were calculated using the Species Demarcation Tool (Muhire et al., 2014) and visualized in RStudio.

### Structural prediction and motif analysis

Protein structures were predicted using AlphaFold3 (Abramson et al., 2024) via the online server (https://alphafoldserver.com/). Predicted structures were converted from CIF to PDB format using the mCIF-to-PDB converter (https://project-gemmi.github.io/wasm/convert/cif2pdb.html). Structural similarity matrices were generated using the DALI server (Holm, 2022). Pairwise structural alignments were performed using the “align” function in PyMOL (https://pymol.org) (DeLano, 2002). Structural similarity was assessed using DALI *z*-scores, with values > 2 considered significant (Holm et al., 2023).

Motif discovery was conducted using the MEME suite (Bailey & Elkan, 1994) with the following parameters: any number of motif repetitions, a maximum of 20 motifs, and motif widths between 10 and 100 amino acids. Motif positions were mapped onto 3D structures and visualized in PyMOL.

### Sequence alignment and phylogenetic analysis

Multiple sequence alignments were performed using MUSCLE v5 with the -super5 option (Edgar, 2022), followed by manual curation. Phylogenetic trees were constructed using the maximum likelihood method implemented in IQ-TREE v2.2.0 (Minh et al., 2020). The best-fit substitution model (LG+F+R5) was selected using ModelFinder (Kalyaanamoorthy et al., 2017). For broader phylogenetic comparisons with cellular organisms, alignments were trimmed to remove poorly aligned regions, and trees were reconstructed using RAxML with the PROTGAMMALG model (Kozlov et al., 2019). All trees were visualized using iTOL v5.0 (https://itol.embl.de).

### Similarity network analysis

To investigate the functional relationships among viral HSP70 proteins and their similarity to cellular homologs, we constructed sequence similarity networks using the Enzyme Function Initiative-Enzyme Similarity Tool (EFI-EST) web server (https://efi.igb.illinois.edu) (Oberg et al., 2023). Viral HSP70 amino acid sequences were submitted to EFI-EST to generate pairwise sequence similarity networks based on BLAST alignment scores. Two alignment score thresholds were applied: a strict threshold of 35 to identify highly similar sequences, and a relaxed threshold of 10 to capture more distant relationships.

To assess relationships with cellular HSP70s, we included representative sequences from plants, animals, fungi, protists, bacteria, and archaea (10 species per group) and generated a network using both relaxed and strict thresholds.

The resulting networks were visualized using Cytoscape v3.9.1 (Shannon et al., 2003).

## Acknowledgments

Computations were performed on the HPC cluster Garnatxa at the I2SysBio. S.F.E was supported by grants PID2022-136912NB-I00 funded by MCIU/AEI/10.13039/501100011033 and by “ERDF a way of making Europe”, and CIPROM/2022/59 funded by Generalitat Valenciana.

## Supplementary data

**Figure S1.** (A) Pairwise amino acid identity among large HSP70 proteins (> 900 amino acids) from *Imitervirales* members: *Acanthamoeba castellanii mimivirus* (BAV61350), *Bandra megavirus* (AUV58293), *Catovirus naegleriensis* (CAK7596984), *Cotonvirus japonicus* (BCS83008), *Moumouvirus australiensis* (AVL94675), and *Yasminevirus* (VBB18370). (B) Pairwise identity among small HSP70 proteins (< 650 amino acids) from *Imitervirales* members: *A. castellanii mimivirus* (BAV61494), *C. naegleriensis* (CAK7597096, CAK7596788), *C. japonicus* (BCS83166), *Megavirus courdo 11* (AFX92555), *M. australiensis* (AVL94812), and *Yasminevirus* (VBB18048).

**Figure S2.** (A) HSP70 gene from *Megavirus courdo 11* showing expression via alternative splicing. (B) Two adjacent HSP70 genes from *M. courdo 11*, separated by an intron-like region. The predicted 3D structures of their translation products align to form a complete HSP70 protein. (C) Fragmented HSP70 genes in *Yasminevirus*, separated by an unrelated intervening ORF. The combined translation products yield a full-length HSP70 protein.

**Figure S3.** Sequence similarity network of viral and cellular HSP70 proteins generated using the EFI-EST tool. (A) Network of the viral HSP70s using a relaxed alignment score threshold of 10. All sequences form a single connected network, but the spatial separation of nodes still reflects two major subgroups, consistent with functional and evolutionary divergence. (C) Network including HSP70s from viruses and representative cellular organisms (10 species each from plants, animals, fungi, protists, bacteria, and archaea), constructed using the relaxed threshold (10). HSP70s from ssRNA viruses form clustered at the edges of the network, while those from dsDNA viruses integrate with cellular HSP70s, mainly from protists.

**Table S1.** List of viruses and their corresponding HSP70 proteins used in this study.

**Table S2.** Number, genomic position, and strand orientation of HSP70 genes in *Imitervirales* genomes.

**Table S3.** Number, genomic position, and strand orientation of HSP70 genes in *Algavirales* genomes.

**Table S4.** List of conserved motifs predicted in viral HSP70 proteins using MEME.

