## Supplementary material for "Multiple origins, one function: evolutionary pathways of HSP70 proteins in viruses": suppl_fig.pdf

**Figure S1**

A

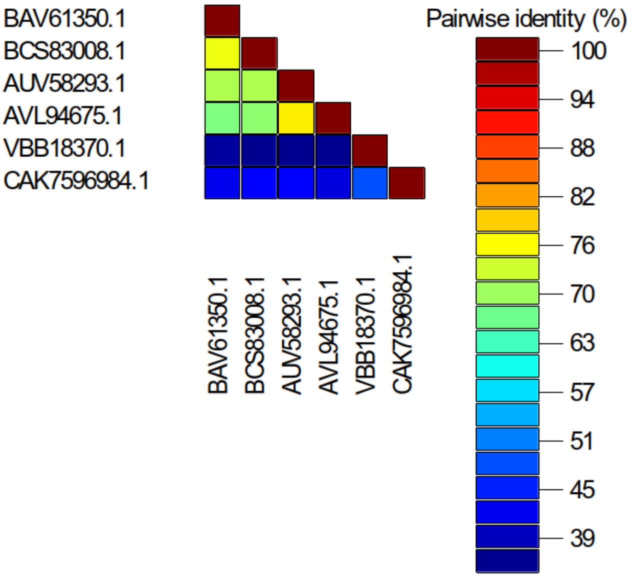


B

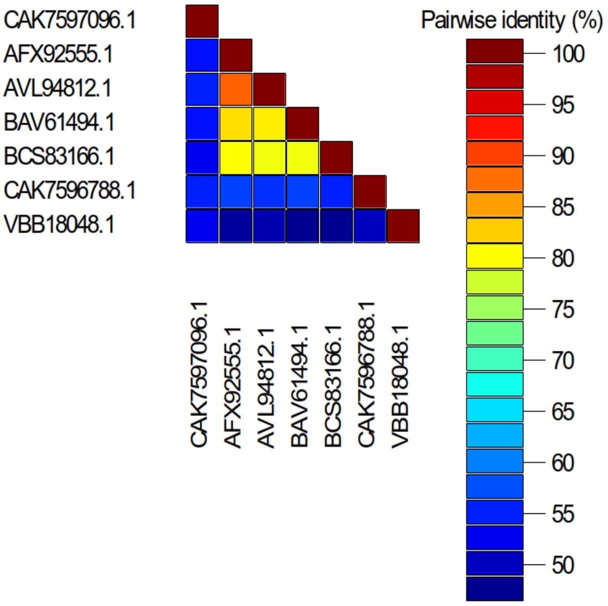


**Figure S1.** A) Pairwise identity between the large heat shock proteins 70 (> 900 amino acids) among Imitervirales members. Bav61350, from acanthamoeba castellani mimivirus; bcs83008, from cotonvirus japonicus; auv58293, bandra megavirus; avl94675, from moumouvirus australiensis; vbb18370, from yasmenivirus; cak7596984, from catovirus naegleriensis. B) Pairwise identity between the small heat shock proteins (<650 amino acids) from the Imitervirales members. Cak7597096 catovirus naegleriensis, ; afx92555, from megavirus courdo 11 ; avl94812, from moumouvirus australiensis ; bav61494, from acanthamoeba castellanii mimivirus; bcs83166, from cotonvirus japonicus; cak7596788, from catovirus naegleriensis; vbb18048, from yasminevirus.

**Figure S2**

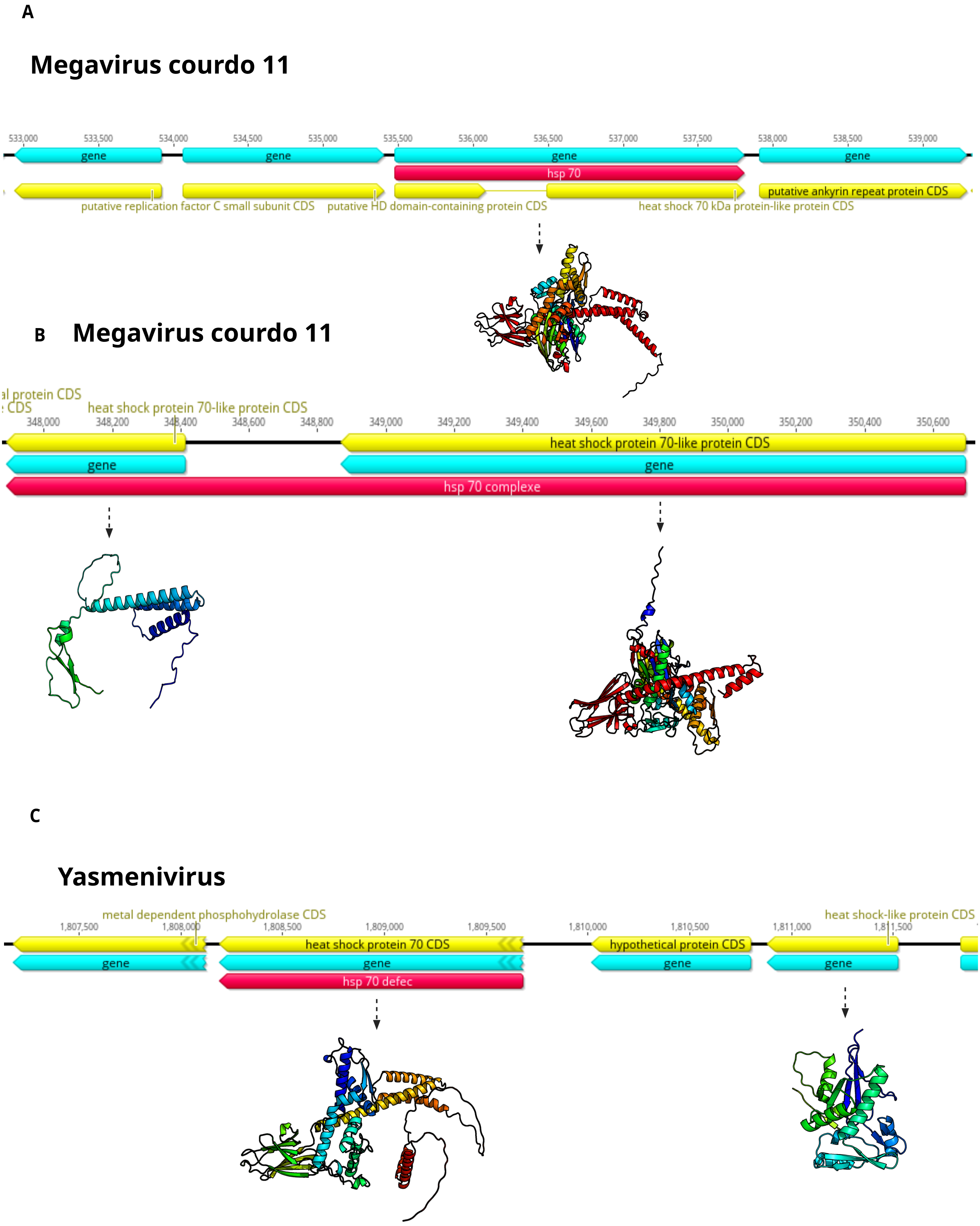


**Figure S2.** Heat shock proteins 70 (HSP70) from megavirus courdo 11, expressed via alternative splicing. B) HSP70 genes from megavirus courdo 11, showing two complete HSP70s split via an intron. The 3D structure of these two proteins yield a complete HSP70. C) HSP70 genes from yasminevirus, showing complete and incomplete HSP70 genes separated by a gene. The 3D structure of these two proteins yield a complete HSP70.

**Figure S3**

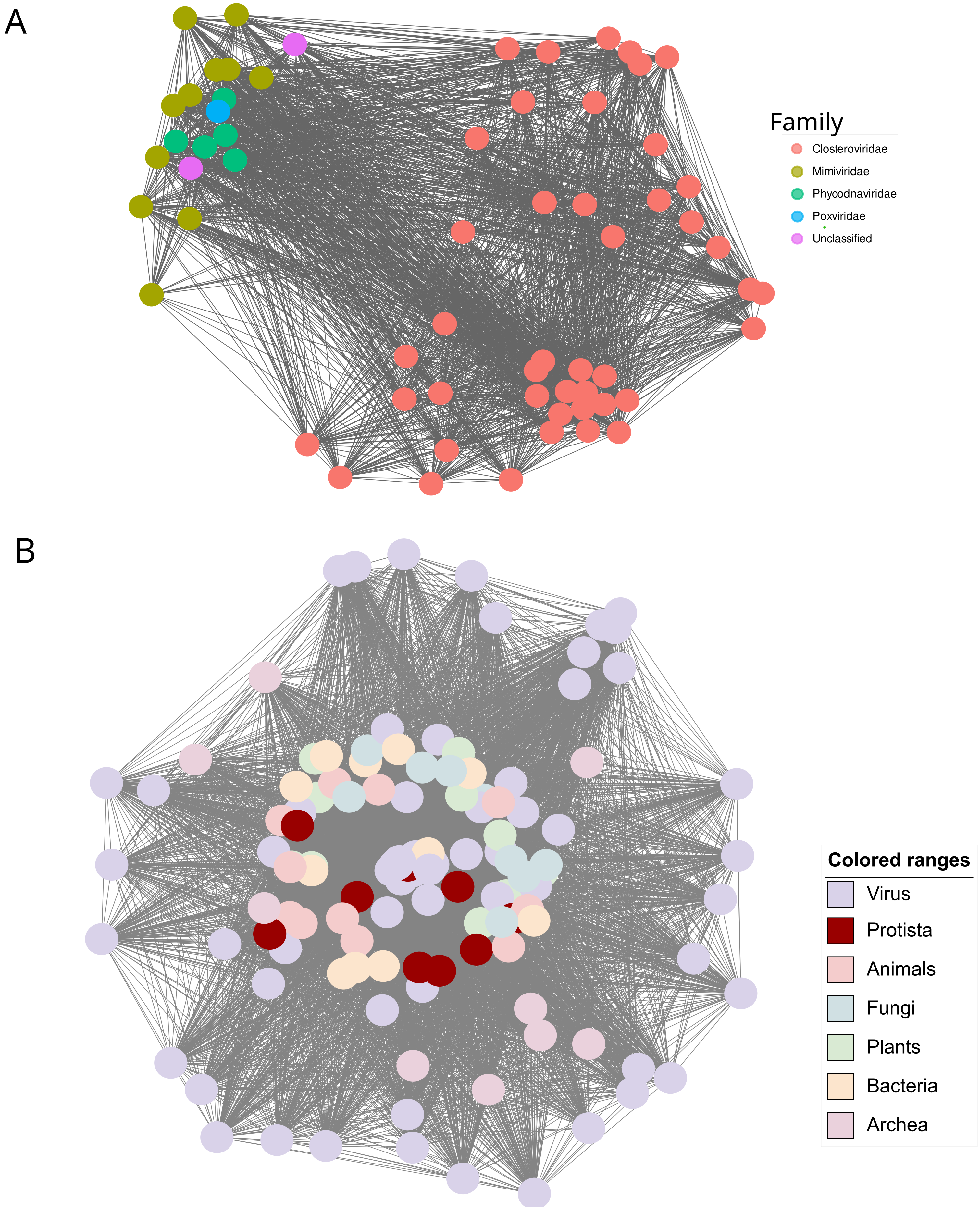


Sequence similarity network of viral and cellular HSP70 proteins generated using the EFI-EST tool. (A) Network of the viral HSP70s using a relaxed alignment score threshold of 10. All sequences form a single connected network, but the spatial separation of nodes still reflects two major subgroups, consistent with functional and evolutionary divergence. (C) Network including HSP70s from viruses and representative cellular organisms (10 species each from plants, animals, fungi, protists, bacteria, and archaea), constructed using the relaxed threshold (10). HSP70s from ssRNA viruses form clustred at the edges of the network, while those from dsDNA viruses integrate with cellular HSP70s, mainly from protists.
