## Supplementary material for "Multiple origins, one function: evolutionary pathways of HSP70 proteins in viruses": TableS4.docx

**Table S4: List of MEME motifs predicted from heat shock protein 70.**

| Motif ID | Motif logo | E-value | Length | Domain^a^ | #of sites^b^ | Comments |
| --- | --- | --- | --- | --- | --- | --- |
| 1 | 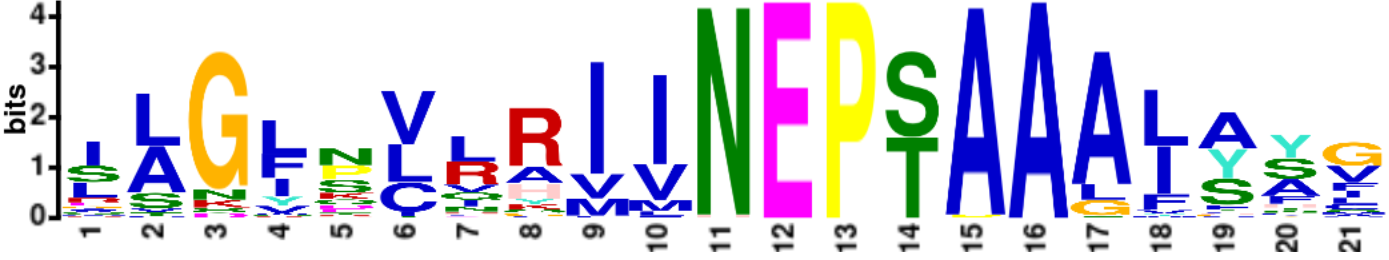 | 9.9e-647 | 21 | NBD | 63 | This motif is present among all viruses |
| 2 | 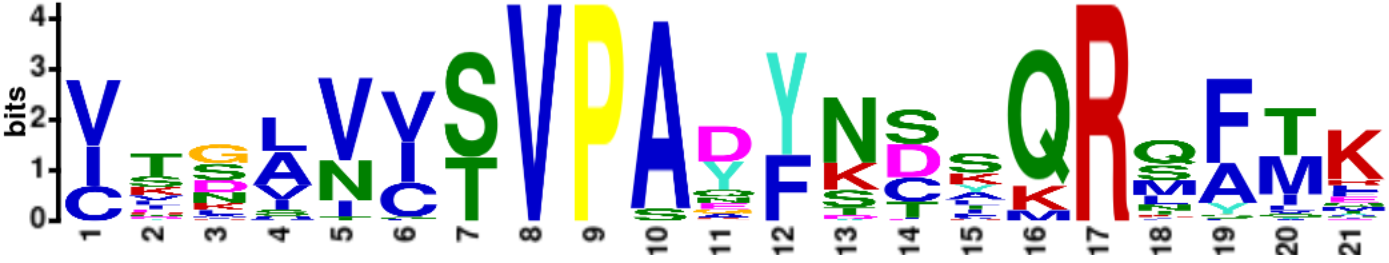 | 7.8e-649 | 21 | NBD | 63 | This motif is present among all viruses |
| 3 | 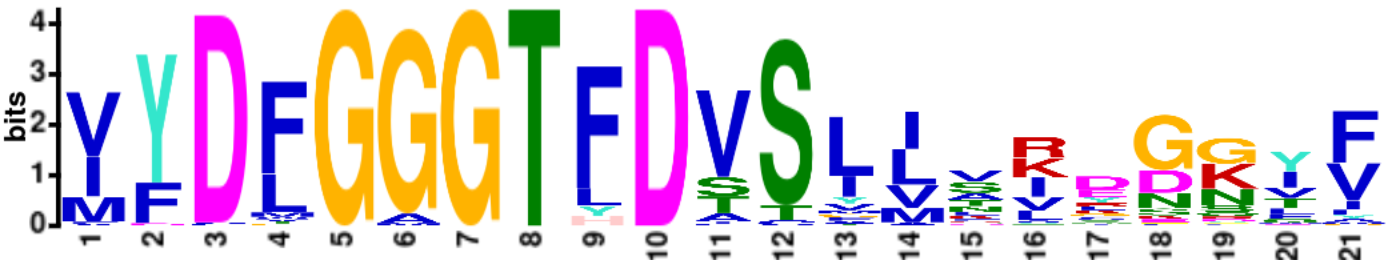 | 5.8e-639 | 21 | NBD | 63 | This motif is present among all viruses |
| 4 | 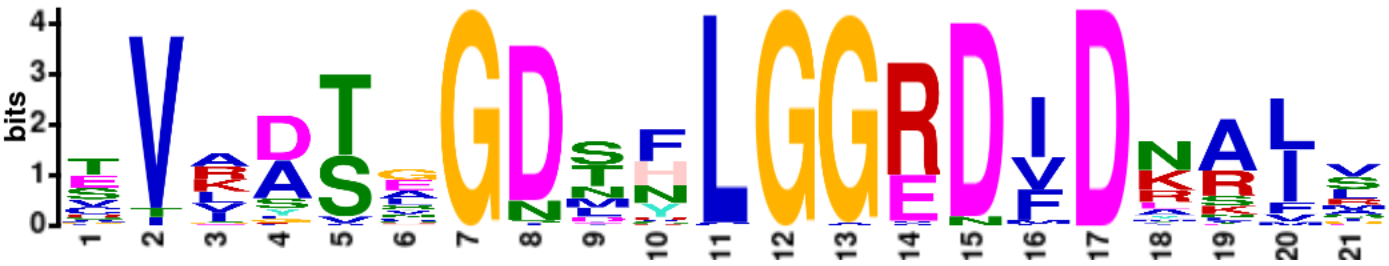 | 1.7e-608 | 21 | NBD | 63 | This motif is present among all viruses |
| 5 | 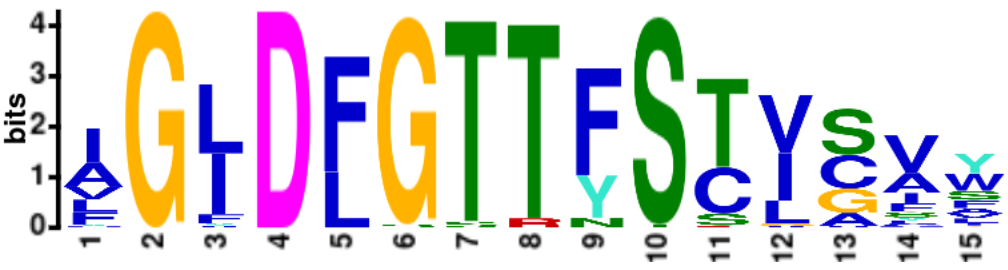 | 3.6e-524 | 15 | NBD | 63 | This motif is present among all viruses |
| 6 | 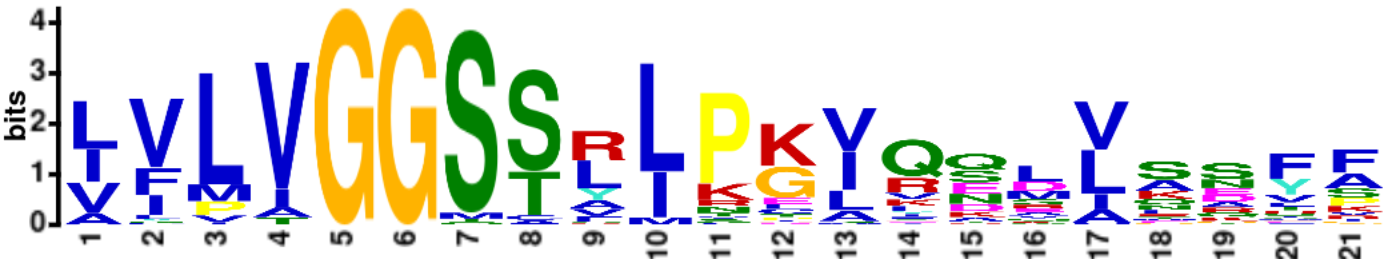 | 1.9e-447 | 21 | NBD | 63 | This motif is present among all viruses |
| 7 | 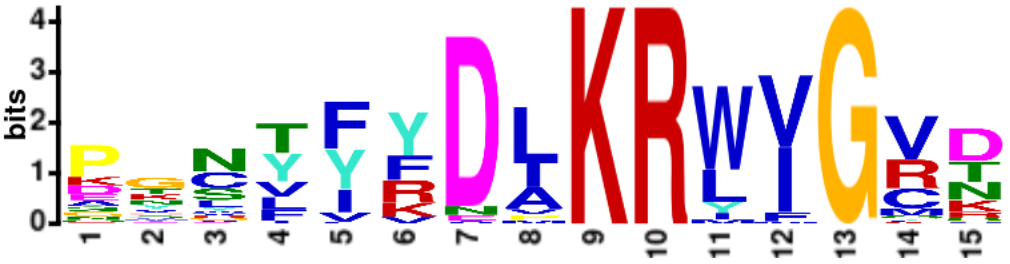 | 2.4e-423 | 15 | NBD | 63 | This motif is present among all viruses |
| 8 | 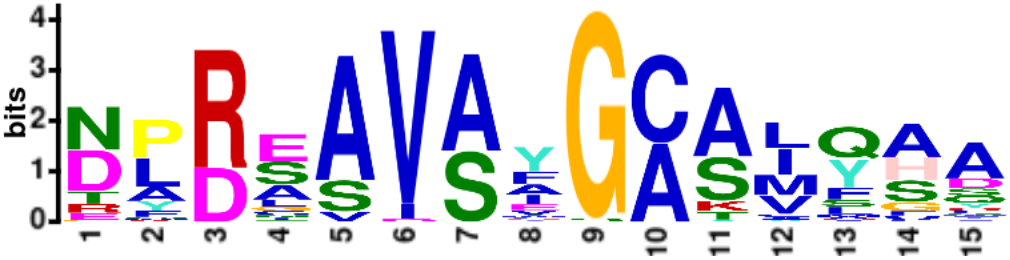 | 1.0e-381 | 15 | NBD | 63 | This motif is present among all viruses |
| 9 | 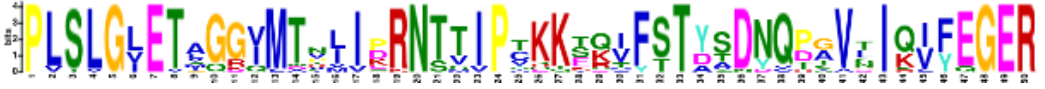 | 8.4e-558 | 50 | SBP | 20 | This motif is present among giant viruses: *Mimiviridae*, *Phycodnaviridae*, Poxviridae, and unclassified ones |
| 10 | 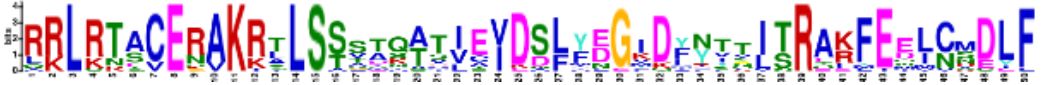 | 4.8e-413 | 50 | NBD | 19 | This motif is present among giant viruses: *Mimiviridae*, *Phycodnaviridae*, and *Poxviridae*. It is absent from Acinetobacter |
| 11 | 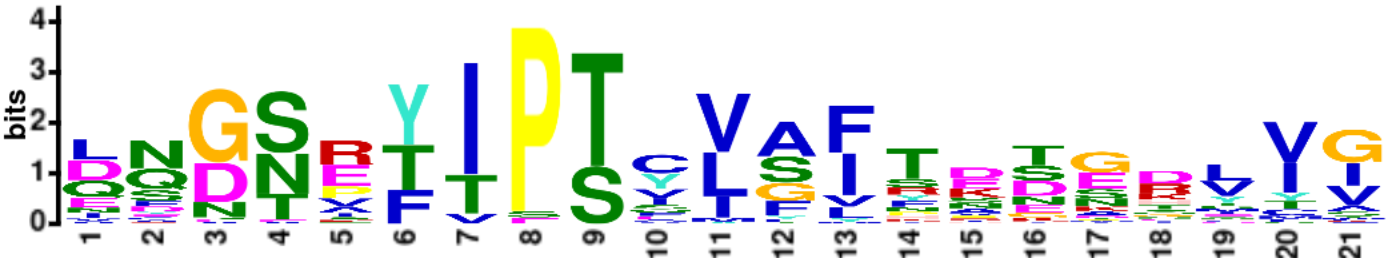 | 1.1e-330 | 21 | NBD | 62 | This motif is present among all viruses except for one virus |
| 12 | 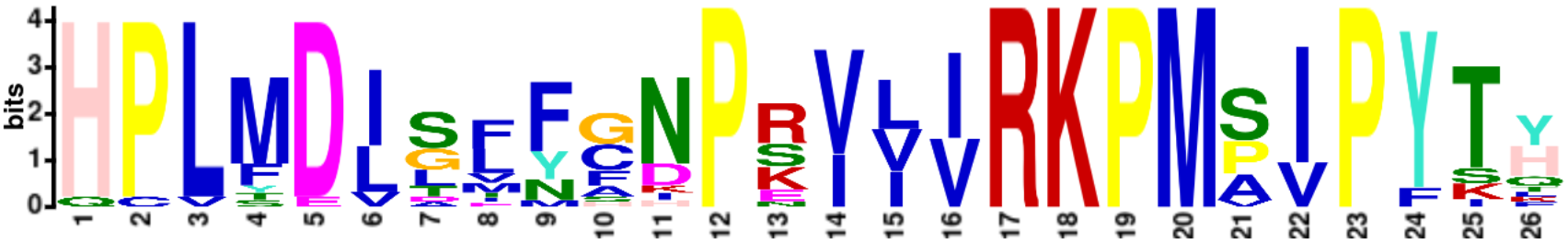 | 2.7e-254 | 26 | SBP | 18 | This motif is present in some viruses belonging to the *Closteroviridae* family: criniviruses, closteroviruses, and velariviruses |
| 13 | 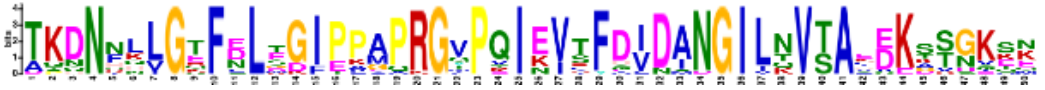 | 2.1e-487 | 50 | SBP | 20 | This motif is present among giant viruses: Mimiviridae, Phycodnaviridae, *Poxviridae*, and unclassified ones |
| 14 | 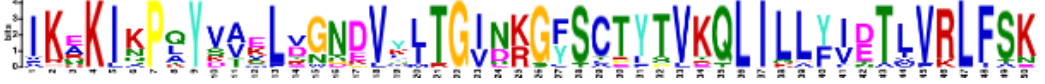 | 3.6e-358 | 50 | NBD | 15 | This motif is present in some viruses belonging to the *Closteroviridae* family: criniviruses, closteroviruses, and menthaviruses |
| 15 | 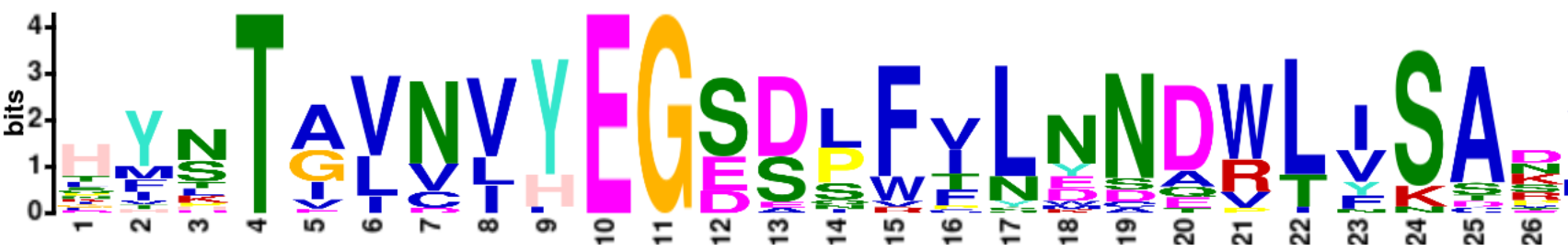 | 4.4e-255 | 29 | SBP | 24 | This motif is present in some viruses belonging to the *Closteroviridae* family: criniviruses, closteroviruses, and velariviruses |
| 16 | 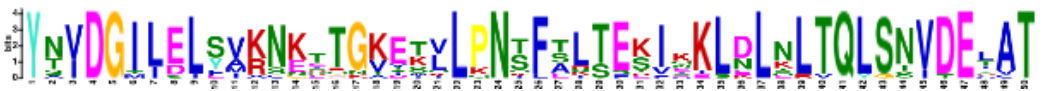 | 8.0e-322 | 50 | NBD | 15 | This motif is present in some viruses belonging to the *Closteroviridae* family: criniviruses, closteroviruses, and velariviruses |
| 17 | 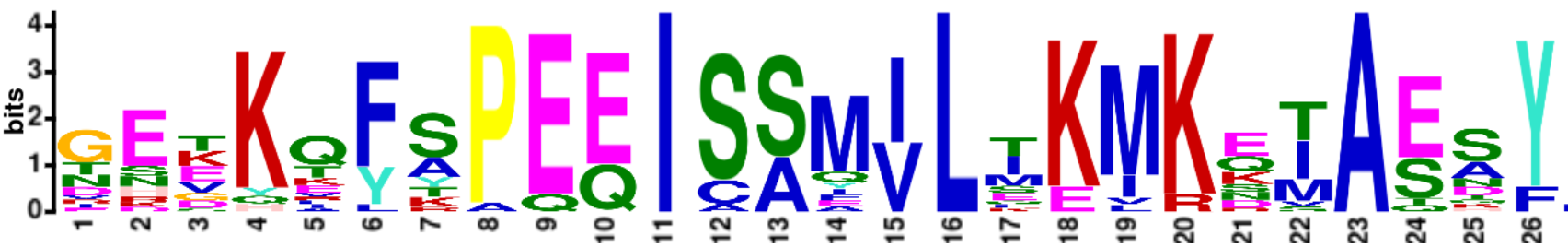 | 2.2e-227 | 29 | NBD | 20 | This motif is present among giant viruses: Mimiviridae, Phycodnaviridae, Poxviridae, and unclassified ones |
| 18 | 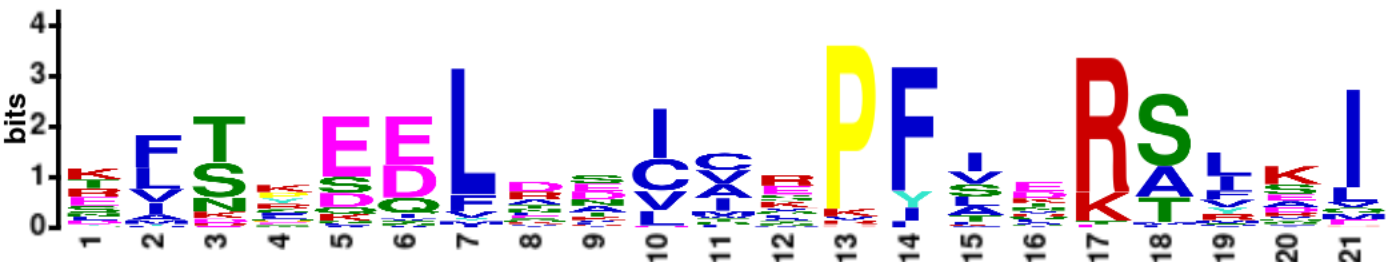 | 4.1e-195 | 21 | SBP | 48 |  |
| 19 | 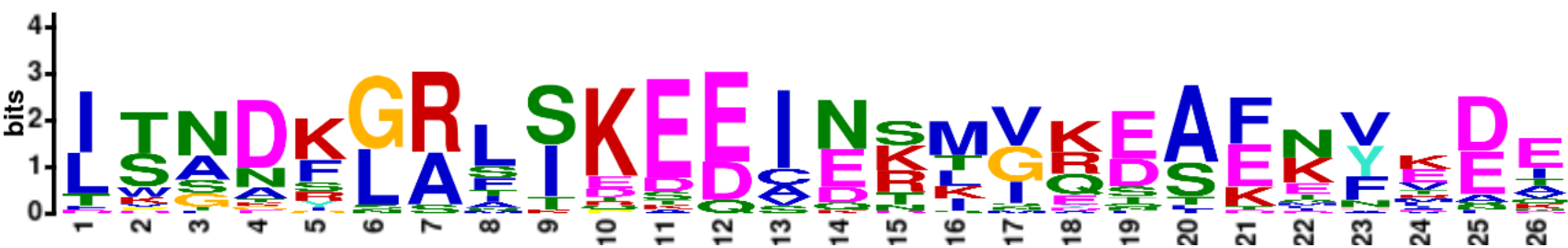 | 3.5e-195 | 29 |  | 30 |  |
| 20 | 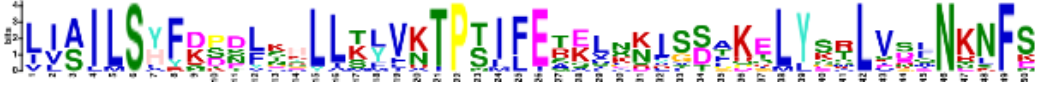 | 1.4e-197 | 50 | Lid | 14 | This motif is present in criniviruses, one closetovirus (plant associated closterovirus 2) |
